# Design of time-delayed safety switches for CRISPR gene therapy

**DOI:** 10.1101/2021.04.17.440285

**Authors:** Dashan Sun

**Affiliations:** Arizona State University, Tempe, Arizona, 85281, US

**Keywords:** ultrasensitivity, time delay, off-target, immune response, CRISPR

## Abstract

CRISPR system is a powerful gene editing tool which has already been reported to address a variety of gene relevant diseases in different cell lines. However, off-target effect and immune response caused by Cas9 remain two fundamental problems. Inspired by previously reported Cas9 self-elimination systems, time-delayed safety switches are designed in this work. Firstly, ultrasensitive relationship is constructed between Cas9-sgRNA (enzyme) and Cas9 plasmids (substrate), which generates the artificial time delay. Then intrinsic time delay in biomolecular activities is revealed by data fitting and utilized in constructing safety switches. The time-delayed safety switches function by separating the gene editing process and self-elimination process, and the tunable delay time may ensure a good balance between gene editing efficiency and side effect minimization. By addressing gene therapy efficiency, off-target effect, immune response and drug accumulation, we hope our safety switches may offer inspiration in realizing safe and efficient gene therapy in humans.

## 1 | Introduction

CRISPR system was found in a bacterial immune system and engineered into a gene editing tool [1]. Specifically, CRISPR system consists of a DNA endonuclease enzyme Cas9 and a single guide RNA (sgRNA) [2]. sgRNA will bind with Cas9, recognize target DNA and localize Cas9 on the target. Because the protospacer element of sgRNA is highly programmable, CRISPR system shows great potential and has been reported to permanently modify disease relevant genes in the liver [3], retina [4], brain [5], heart [6], and skeletal muscle [7]. One of the challenges for the use of CRISPR system for both research and therapeutic application is the off-target effect. To address this problem, both Cas9 [8] and sgRNA [9-10] can be modified to avoid double strand break. However, bacterially derived Cas9 still remains in edited cells and permanent Cas9 expression may induce immune response [11-12]. To cease the permanent Cas9 expression, Cas9 self-deleting systems are developed by introducing a sgRNA which targets the Cas9 plasmids. For *in vitro* experiments [13-15], off-target effects are reported to be obviously decreased while on-target efficiency basically remains the same, and Cas9 protein can be cleared. However, *in vivo* studies reveal that Cas9 self-elimination will decrease on-target efficiency either significantly [16] or slightly [17]. A simple self-targeting design naturally generates a linear pattern of Cas9 self-elimination, and this linear pattern of self-elimination can be too fast for some of the clinical cases and thereby gene therapy efficiency will be decreased.

Ideally, we may expect the Cas9 to remain a certain amount for a high gene therapy efficiency; after gene therapy is complete, Cas9 should be eliminated as fast as possible to minimize side effects. In another word, what we expect is a time delay between gene therapy and Cas9 elimination. In this paper, an ultrasensitivity transmission module is designed to generate the desired time delay. We proved that the time-delayed safety switch is superior to the linear pattern safety switch in one gene therapy case, and we demonstrate that the length of delay time is highly tunable for different gene therapy cases based on mathematical modeling results. Apart from the artificial time delay generated by ultrasensitivity, our data fitting work revealed the intrinsic time delay in biomolecular activities. Safety switches utilizing the intrinsic time delay are designed and tested with fitted parameters. Featured by a time-delayed design, our safety switches have the potential to guarantee gene therapy efficiency when minimizing both off-target and immune response risk, thereby enable safe and efficient gene therapy in humans.

## 2 | Materials and methods

We presented mathematical modeling of ultrasensitivity transmission module, time-delayed safety switches, Cas9 activity and Cre recombination with ordinary differential equations (ODEs) and delayed differential equations (DDEs). The lsqcurvefit function is used for data fitting. Law of mass action for biochemical reactions was used. Reactions are described as equation 1-26; parameters are described as Table 1. Differential equations were solved using the ODE45 and DDE23 routine of MatlabR2020b.

**TA B L E 1.**
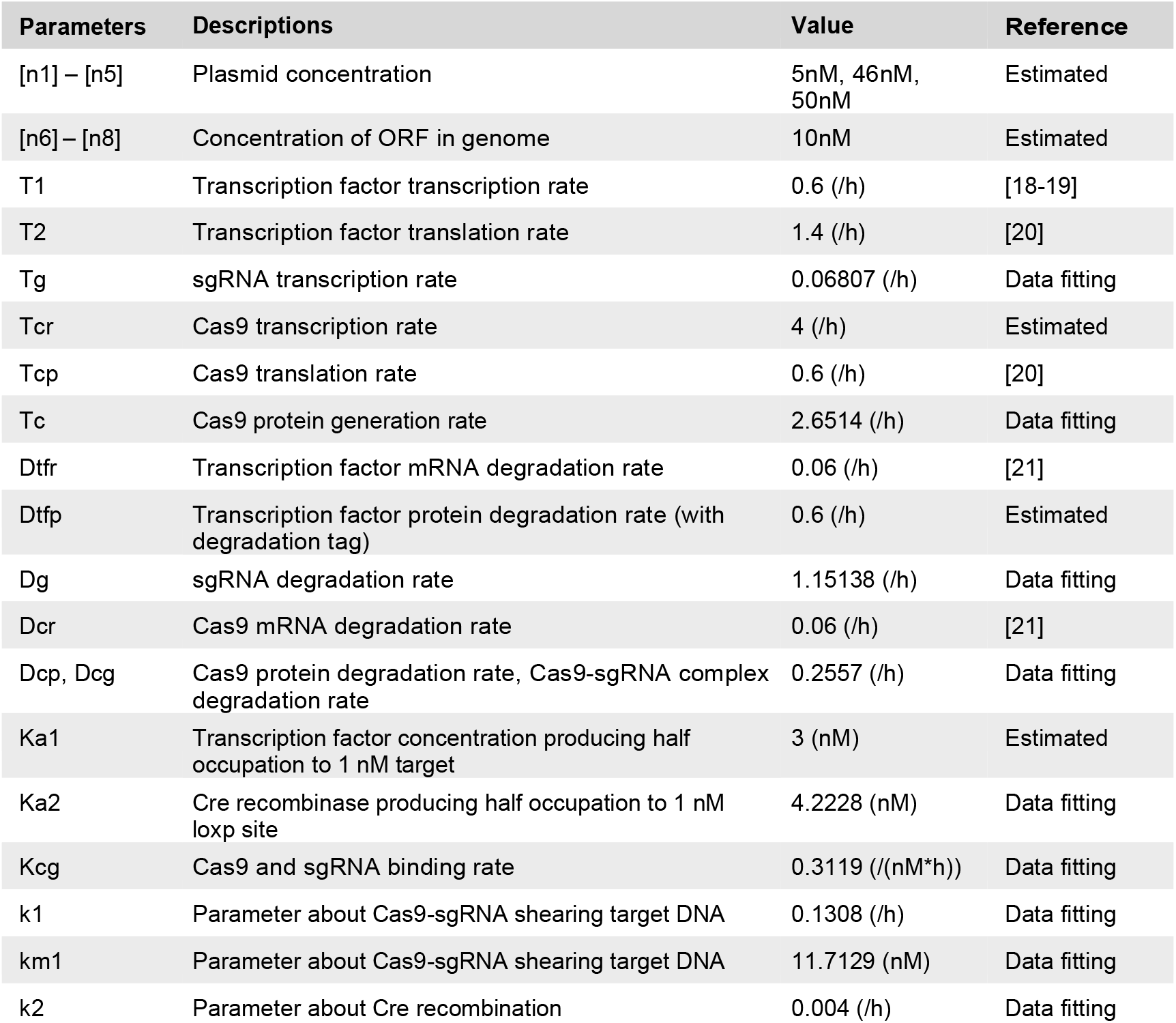
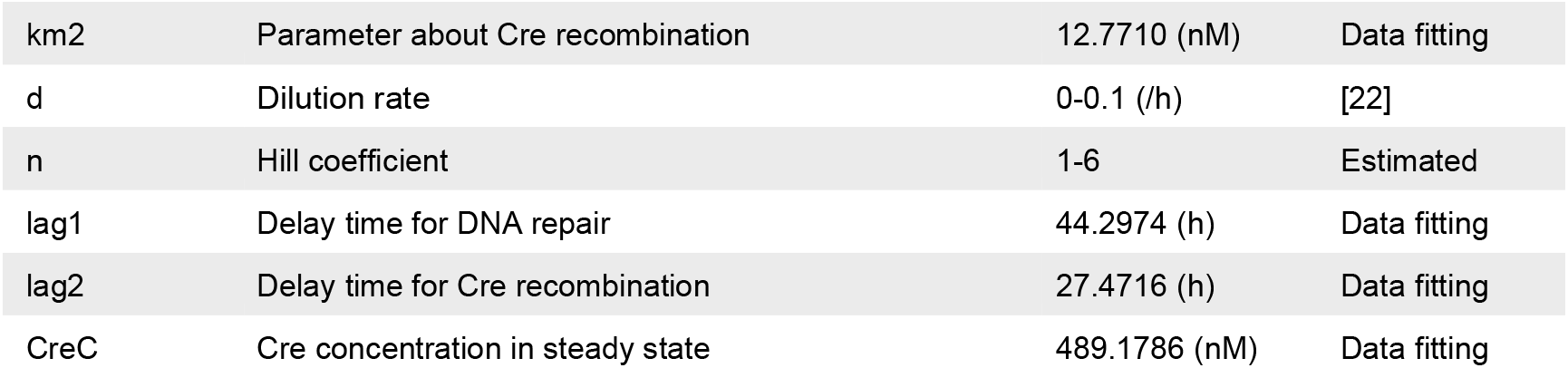
Parameters

## 3 | Results

### 3.1 | Artificial time delay in Cas9 elimination

Previously reported self-deleting CRISPR system [13-17] sets a good example for the design of safety switches and inspires us a lot. In that system, the speed of Cas9 self-deletion is reported to be in a linear fashion. As a certain concentration of Cas9 plasmid is necessary for efficient gene therapy, the linear self-deletion pattern may either require a high initial concentration of Cas9 plasmid and thereby induce immune response; or fail the gene therapy purpose because of the insufficiency of Cas9. Thus, we may expect the speed of self-deletion to be in a non-linear fashion: in the process of gene therapy, the speed of Cas9 self-deletion is slow, thereby gene therapy efficiency can be ensured by a much smaller Cas9 dose; after gene therapy is done, the speed of Cas9 self-deletion is switched to high to minimize side effects. To fulfill this expectation, a time-delayed Cas9 elimination system is designed (Fig. 1A-B). Ultrasensitive relationship between transcription factor and promoter (Fig.1C) is utilized as the fundamental mechanism for generating the time-delayed feature. By introducing Cas9 to the system, the ultrasensitive response to transcription factor is transmitted from sgRNA to Cas9-sgRNA complex. When transcription factor promoter and Cas9 promoter are designed to have the same sgRNA target site, ultrasensitive relationship between Cas9-sgRNA (enzyme) and Cas9 plasmid (substrate) is established (Fig. 1D). Based on this ultrasensitivity transmission module, time-delayed Cas9 elimination is realized and the switch from slow Cas9 elimination to rapid Cas9 elimination is automatic.

**FIGURE 1.**
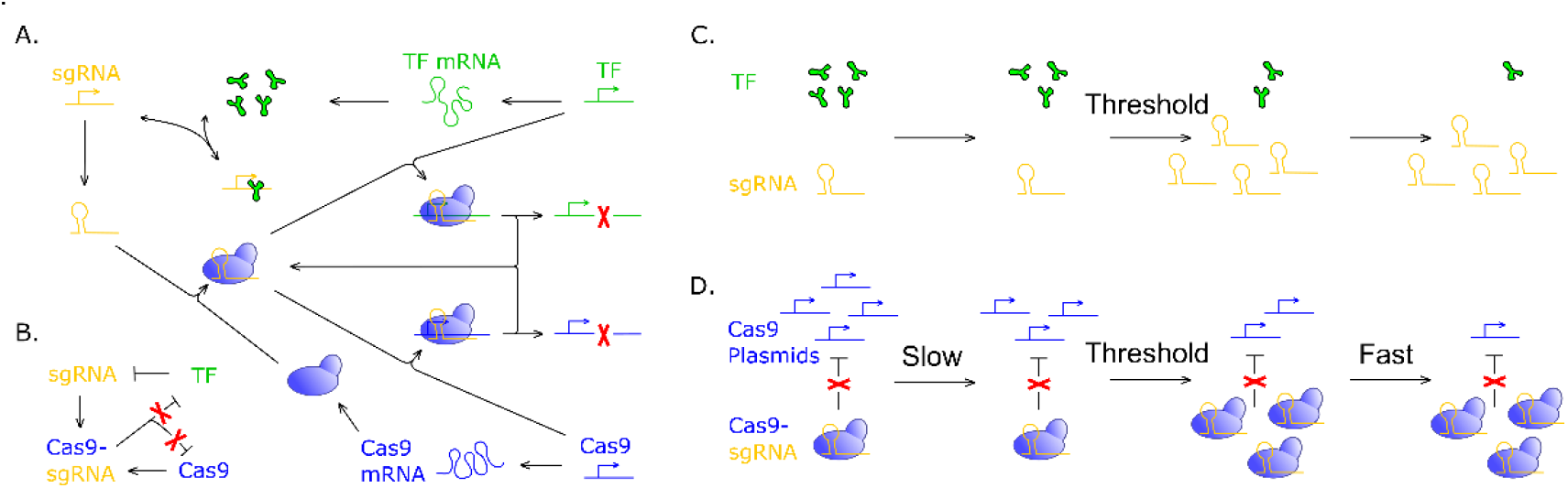
Time-delayed Cas9 elimination system. TF stands for transcription factor. (A) Detailed molecular mechanism of time-delayed Cas9 elimination system. (B) Kinetic schemes for time-delayed Cas9 elimination system. (C) Ultrasensitive relationship between transcription factor and sgRNA transcription. sgRNA transcription rate changes dramatically around a threshold concentration of transcription factor. (D) Ultrasensitive relationship between Cas9 plasmid (substrate) and Cas9-sgRNA (enzyme) is developed by circuit connection, which yields time-delayed Cas9 elimination.

#### 3.1.1 | Ultrasensitivity and time delay

Ultrasensitivity describes a relationship between output and stimulus which is more sensitive than the hyperbolic Michaelis-Menten response. Ultrasensitivity is capable of filtering out noise and generating fast activation around threshold [23]; and plays important roles in a variety of cellular events like cell fate determination [24]. A broad range of methods can be used for achieving ultrasensitivity, such as multistep [25], saturation mechanism [26] and positive feedback [27]. In our work, cooperativity of transcription factors binding the target site is utilized for achieving ultrasensitivity. Firstly, a transcription factor is employed to repress the promoter of sgRNA (Fig. 2A). mRNA of transcription factor is transcribed under a constitutive promoter (equation 1), then transcription factor protein is translated (equation 2) and repress the sgRNA transcription (equation 3).

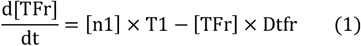

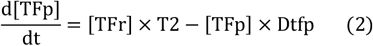

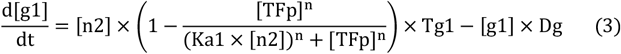

**FIGURE 2.**
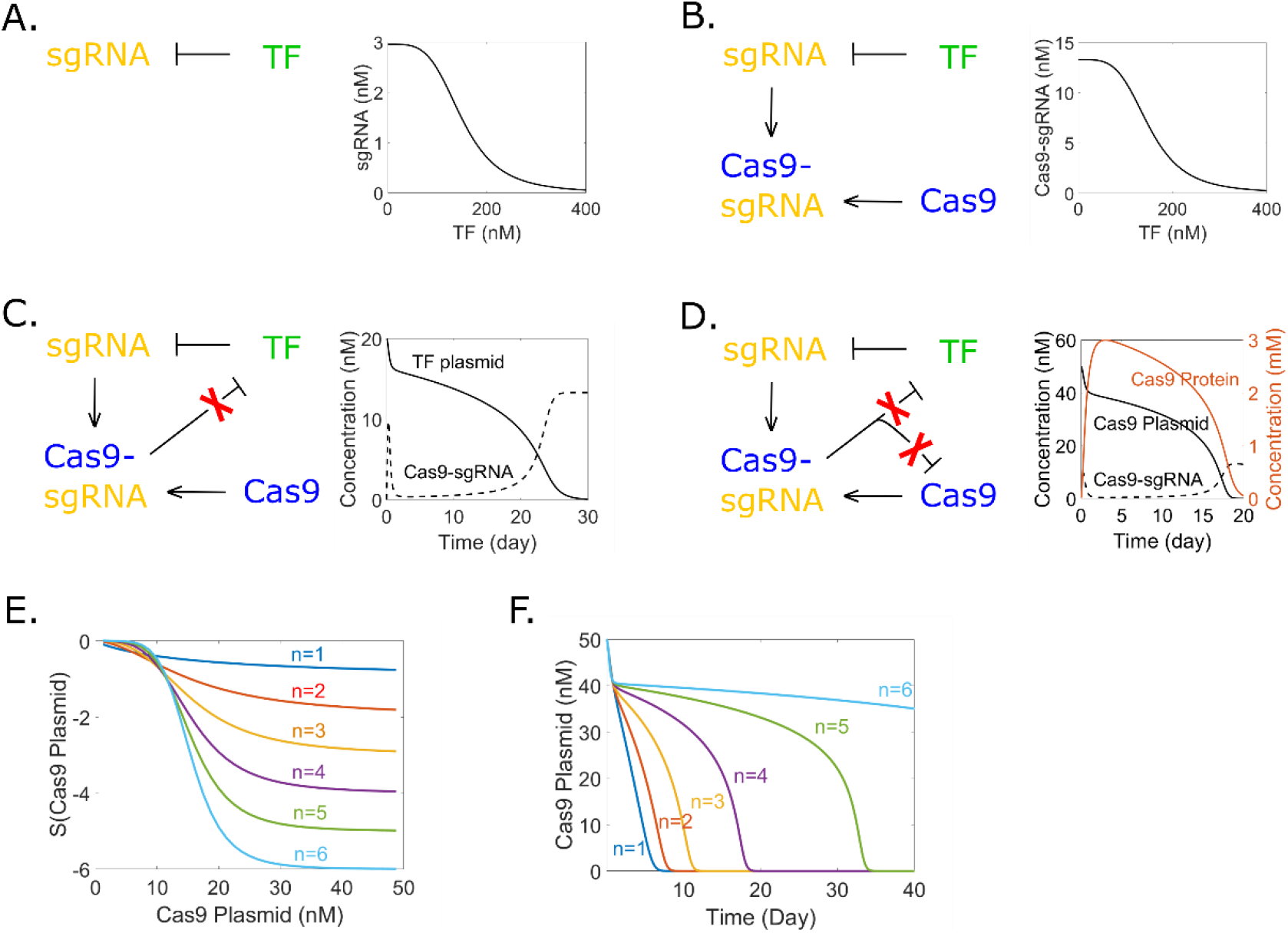
Ultrasensitivity transmission and time delay. TF stands for transcription factor. (A) Cooperativity of transcription factor binding the target site generates the ultrasensitivity. sgRNA concentration (output) is highly sensitive to TF concentration (input) around a threshold. (B) Ultrasensitive relationship between Cas9-sgRNA and TF. (C) Time delayed TF elimination. Ultrasensitive relationship is built between Cas9-sgRNA (enzyme) and TF (substrate). (D) Time-delayed Cas9 elimination. Ultrasensitive relationship is built between Cas9-sgRNA (enzyme) and Cas9 plasmid (substrate). Response of Cas9 protein has a short delay compared to Cas9 plasmid. Cas9 protein maintains a certain value before rapid elimination. (E) Response of Cas9-sgRNA concentration (output) to Cas9 plasmid concentration (stimulus) is more sensitive with higher Hill coefficient n (n=1, 2, 3, 4, 5, 6). (F) Higher value of Hill coefficient n leads to longer delay time and larger difference in Cas9 elimination speed.

Where [TFr], [TFp] and [g1] are defined as transcription factor mRNA concentration, transcription factor protein concentration and sgRNA concentration, respectively. T1, T2 and Tg1 stands for transcription factor transcription rate, translation rate and sgRNA transcription rate, respectively. Dtfr, Dtfp and Dg stands for degradation rates. Plasmid concentration is defined as n1 and n2. Ka1 is the ratio of transcription factor to target which produces half occupation and n is the Hill coefficient. (Table 1)

To show the ultrasensitive relationship between sgRNA and transcription factor, steady state response curve is given (Fig. 2A) by solving equations 1-3, whereas sgRNA concentration is the output and transcription factor concentration is the stimulus. To quantify the sensitivity and further confirm ultrasensitivity, a sensitivity analysis is performed, and the result (Fig. S1) proves the ultrasensitive relationship.

After parameters are set to an appropriate value which enable the transcription factor to repress sgRNA well, Cas9 is introduced to the system (equation 4-5). Equation 3 is revised as equation 6 as Cas9 will bind with sgRNA and form the Cas9-sgRNA complex (equation 7).

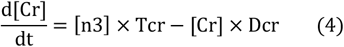

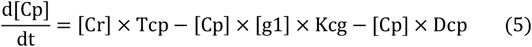

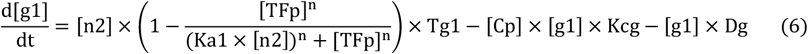

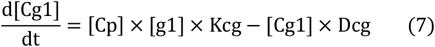

Where [Cr], [Cp] and [Cg1] are defined as Cas9 mRNA concentration, Cas9 protein concentration and Cas9-sgRNA complex concentration, respectively. [n3] is defined as Cas9 plasmid concentration. Tcr stands for Cas9 transcription rate, Tcp stands for Cas9 translation rate, Kcg stands for Cas9 and sgRNA binding rate; Dcr, Dcp and Dcg are degradation rates. (Table 1)

By solving equations 1-2, 4-7, the steady state response curve is given as Fig. 2B. A further sensitivity analysis (Fig. S2) proves the ultrasensitive relationship between Cas9-sgRNA and TF. From the steady state response curve we can see that, a high initial concentration of TF plasmids gives a low concentration of Cas9-sgRNA. If sgRNA is designed to target TF plasmids (equation 8), TF plasmids concentration will gradually decrease. Before TF concentration decrease to the threshold, the Cas9-sgRNA concentration remains low. Considering that Cas9-sgRNA shears TF plasmids like an enzyme, the speed of TF elimination should be slow in the beginning. Once TF concentration decrease to the threshold, there will be a rapid increase in Cas9-sgRNA concentration because of the ultrasensitivity. The speed of TF elimination will be boomed as a result, which will further accelerate Cas9-sgRNA accumulation and form a positive feedback. Therefore, we expect to see a significant change in the speed of TF elimination around the threshold.

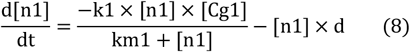

Where k1 and km1 are parameters describing the activity of enzyme catalyzing substrates; d is the dilution rate which describes the speed of cell division. d is set to 0 when trying to get steady states of the system. (Table 1)

The result of solving equations 1-2, 4-8 (Fig. 2C) fulfills our expectation. leakage of sgRNA exists in the beginning because it takes some time for transcription factor to be transcribed and translated after transfection. Because sgRNA is repressed for about 20 days, Cas9-sgRNA complex stays in low concentration and eliminates TF plasmids slowly in this period. After TF plasmid concentration drops to the threshold (around 10nM), a dramatic increase of Cas9-sgRNA complex can be seen because sgRNA transcription is reactivated. The speed of TF elimination after 20day is about 3 times higher than the speed of TF elimination before 20day. The time delayed TF elimination then needs to be transformed into time delayed Cas9 elimination, and synthetic biology makes it possible to insert the same sgRNA target site into Cas9 plasmids (equation 9-10).

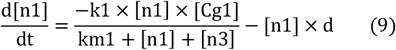

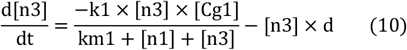

By solving equations 1-2, 4-7 and 9-10, the time delayed Cas9 elimination is obtained as Fig. 2D. Ultrasensitive relationship between Cas9-sgRNA and Cas9 plasmids is generated by inserting sgRNA target site into both TF and Cas9 plasmids. The speed of Cas9 elimination after 15day is about 8 times higher than the speed of TF elimination before 15day, and 15day is the delay time. Response curve of Cas9 protein has a short delay compared to Cas9 plasmids and also has a time-delayed feature for dynamic process. Cas9 will be cleared around day 20.

Given that the time-delayed feature is closely related to the ultrasensitive relationship between transcription factor and sgRNA transcription, we further studied how different kinds of transcription factor (different Hill coefficient n) will affect the time-delayed feature. Sensitivity analysis (see supporting information for introduction and method) for different Hill coefficient n (n=1, 2, 3, 4, 5, 6) is performed and the result (Fig. 2E) indicates that a higher Hill coefficient will increase the sensitivity of Cas9-sgRNA generation (output) to Cas9 plasmid concentration (stimulus). Higher sensitivity will increase the length of delay time and difference in Cas9 elimination speed in the time-delayed Cas9 elimination system (Fig. 2F).

#### 3.1.2 | Gene therapy with time-delayed system elimination

We then compared the time-delayed self-elimination system and linear self-elimination system in a same gene therapy case. In the pursue of a clean self-elimination and thereby minimal drug accumulation, we hope that sgRNA can be eliminated as well. The primary attempt of making sgRNA to also eliminate itself seems simple and direct, but the system will never switch to a fast self-elimination state because the ratio of TF to sgRNA plasmids will be constant and thereby never drop to the threshold. In order to eliminate TF faster than sgRNA plasmids, another sgRNA (sgRNA2) is introduced to the system as Fig. 3A. sgRNA3 targets the genome and serves for the therapy purpose. The time-delayed self-elimination system is modeled with equation 1-2, 4, 7, 11-22. The linear self-elimination system (Fig. 3B) can be modeled with the same equations by simply setting n1 and n4 to 0.

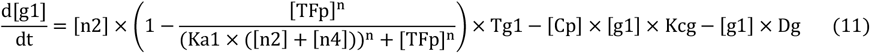

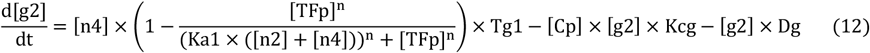

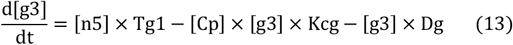

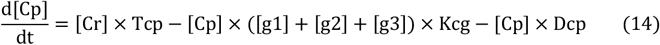

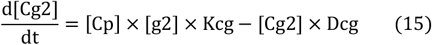

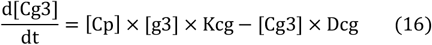

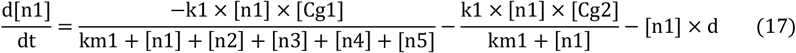

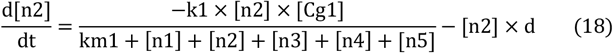

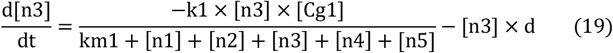

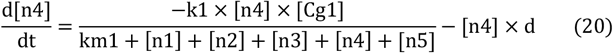

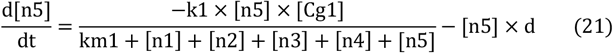

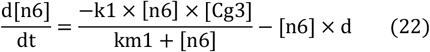

**FIGURE 3.**
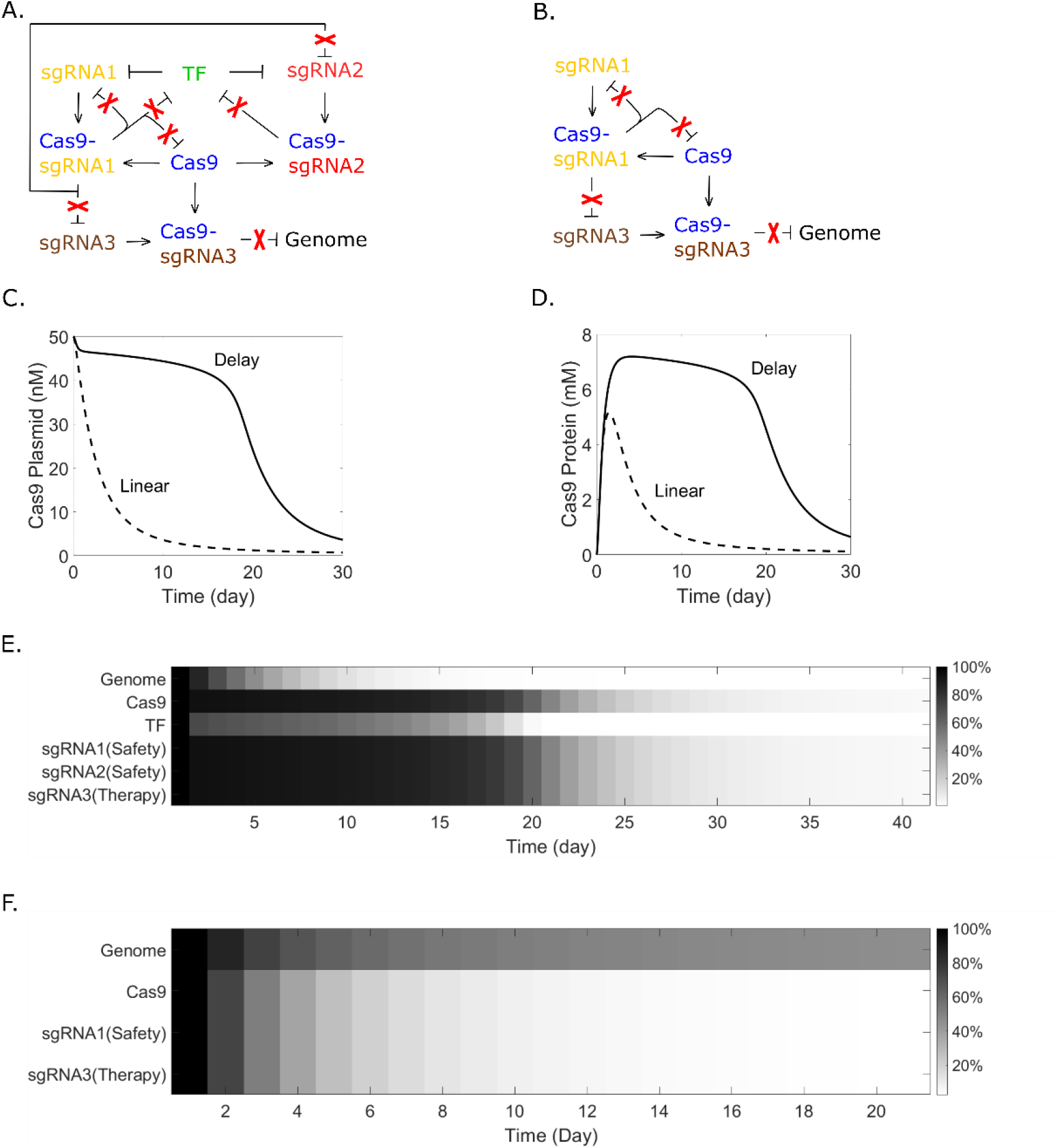
Compare time-delayed self-elimination and linear self-elimination. (A) Time-delayed self-elimination system. sgRNA1 serves for safety purpose, sgRNA2 helps generating the time delay, sgRNA3 serves for therapy purpose. (B) Linear self-elimination system. (C) Dynamic process of Cas9 plasmid elimination. Time-delayed self-elimination system maintains a high Cas9 plasmid concentration before rapid elimination. (D) Dynamic process of Cas9 protein elimination. Cas9 protein maintains a high concentration before rapid elimination. (E) Dynamic process of time-delayed system self-elimination. Gene therapy efficiency is ensured, all components are cleared finally. (F) Dynamic process of linear system self-elimination. A linear self-elimination system may have low gene therapy efficiency.

Where [g2], [g3], [Cg2] and [Cg3] are defined as sgRNA2 concentration, sgRNA3 concentration, Cas9-sgRNA2 complex concentration and Cas9-sgRNA3 complex concentration, respectively. [n1] - [n5] are plasmid concentrations, [n6] refers to genome. (Table 1)

The dynamic process of Cas9 plasmid elimination is illustrated as Fig. 3C. Compared to the linear self-elimination system which may fail the gene therapy purpose even with a high initial Cas9 plasmid dose, the time-delayed self-elimination system is capable of maintain sufficient Cas9 for a while before fast elimination (Fig.3D). For a gene therapy case which takes about 12 days, time-delayed self-elimination system ensures both gene therapy efficiency and safety (Fig. 3E), while linear self-elimination may not complete gene therapy (Fig. 3F).

#### 3.1.3 | Tunable delay time for specific clinical cases

Because of the diversity of diseases and cell lines, and the variety of off-target sites and frequency, delay time which gives best consideration to both gene therapy efficiency and side effects may be different for specific clinical cases. Here we demonstrate that the delay time in our self-elimination system is highly tunable. Taking parameters relevant with TF as an example, by using different promoters (different T1) for TF or changing initial TF plasmid dose (different n1), we can see from Fig. 4 that the delay time is highly tunable from several days to more than 1 moth. Delay time is defined as the time point when Cas9 plasmid concentration drops to threshold concentration and induces maximum sgRNA response.

**FIGURE 4.**
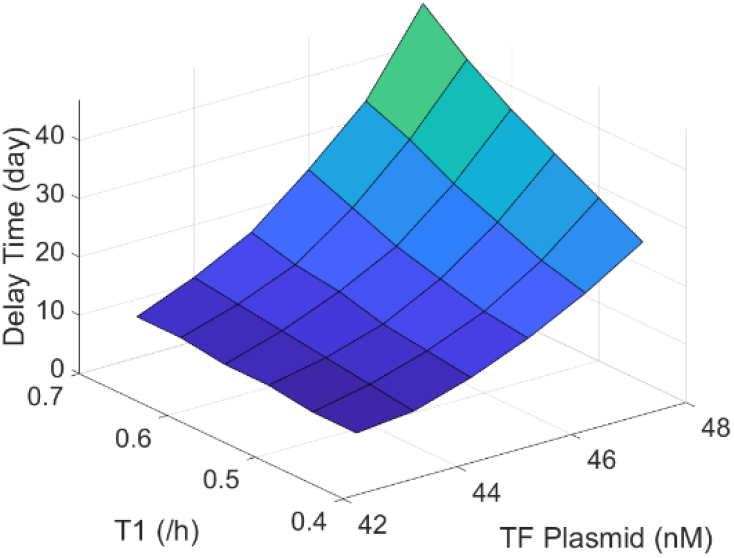
Tunable delay time. Delay time is highly tunable with different promoter for TF (T1 = 0.45/h, 0.5/h, 0.55/h, 0.6/h, 0.65/h, 0.7/h) or plasmid dose (n1 = 43nM, 44nM, 45nM, 46nM, 47nM, 48nM)

#### 3.1.4 | cell division and self-elimination

Dilution rate d is an important parameter which describes cell division rate. However, when trying to get steady state curves for understanding mechanism and feature of the time-delayed self-elimination system, dilution rate has to be set to 0. Thus, it is necessary to see how dilution rate will affect the function of time-delayed self -elimination system. The result of solving equation 1-2, 4, 7, 11-22 with different dilution rate is shown as Fig. 5.

**FIGURE 5.**
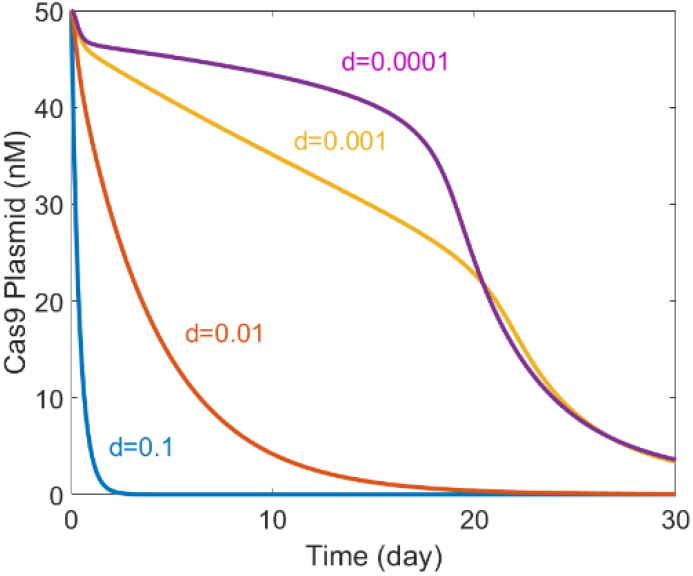
Time-delayed Cas9 elimination in different cell lines (d=0.1, 0.01, 0.001, 0.0001). d stands for dilution rate.

When dilution rate is included in the system, there are two factors responsible for Cas9 plasmid elimination. One factor is cell division and apoptosis, another factor is the self-elimination activity of the system. From Fig. 5 we can see that, when dilution rate is high (d=0.1, 0.01), cell division and apoptosis will play a leading role in Cas9 plasmid elimination, thereby the activity of self-elimination system is not obvious. When the dilution rate is low (d=0.001, 0.0001), the time-delayed self-elimination system will play the leading role in Cas9 elimination. Thus, for cell lines whose division rate is high, the necessity of self-elimination system may need to be double checked. While for cell lines like cardiac cells whose division rate is low, a self-elimination system for CRISPR gene therapy is desired for sure.

### 3.2 | Intrinsic time delay in biomolecular activities

Apart from the artificial time delay generated by ultrasensitivity, Intrinsic time delay in biomolecular activities can also be adopted to construct safety switches for CRISPR gene therapy. Our data fitting work revealed the intrinsic time delay in both DNA repair process and Cre recombination. Safety switches are then designed based on intrinsic time delay and tested with fitted parameters.

#### 3.2.1 | Model construction and data fitting

To address the off-target effect, both Cas9 and sgRNA can be modified to avoid double strand break. From the work carried out in a world-wide scale, when CRISPR does not induce double strand break, we can always observe an immediate CRISPR activity [22,28-29] after transfection. In contrast, when double strand break exists, mutation rate detected by next generation sequencing always have a delay for several days after transfection [16,30]. Based on these observations, we suspect that there can be a time delay in CRISPR induced DNA repair. Among all these works, Krzysztof et al. [30] provides valuable data which reveals the dynamic process and helps the work of system biology a lot. In one of the circuits (Fig. 6A) they tested, Cas9-sgRNA complex induces double strand break in EGFP and mutation rate is detected at different time points. To test the assumption that there’s a time delay in CRISPR induced DNA repair, we constructed two models which differs from each other in the assumption of delayed DNA repair (equation 23) or immediate DNA repair (equation 24).

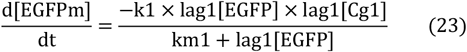

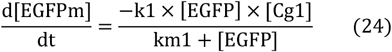

**FIGURE 6.**
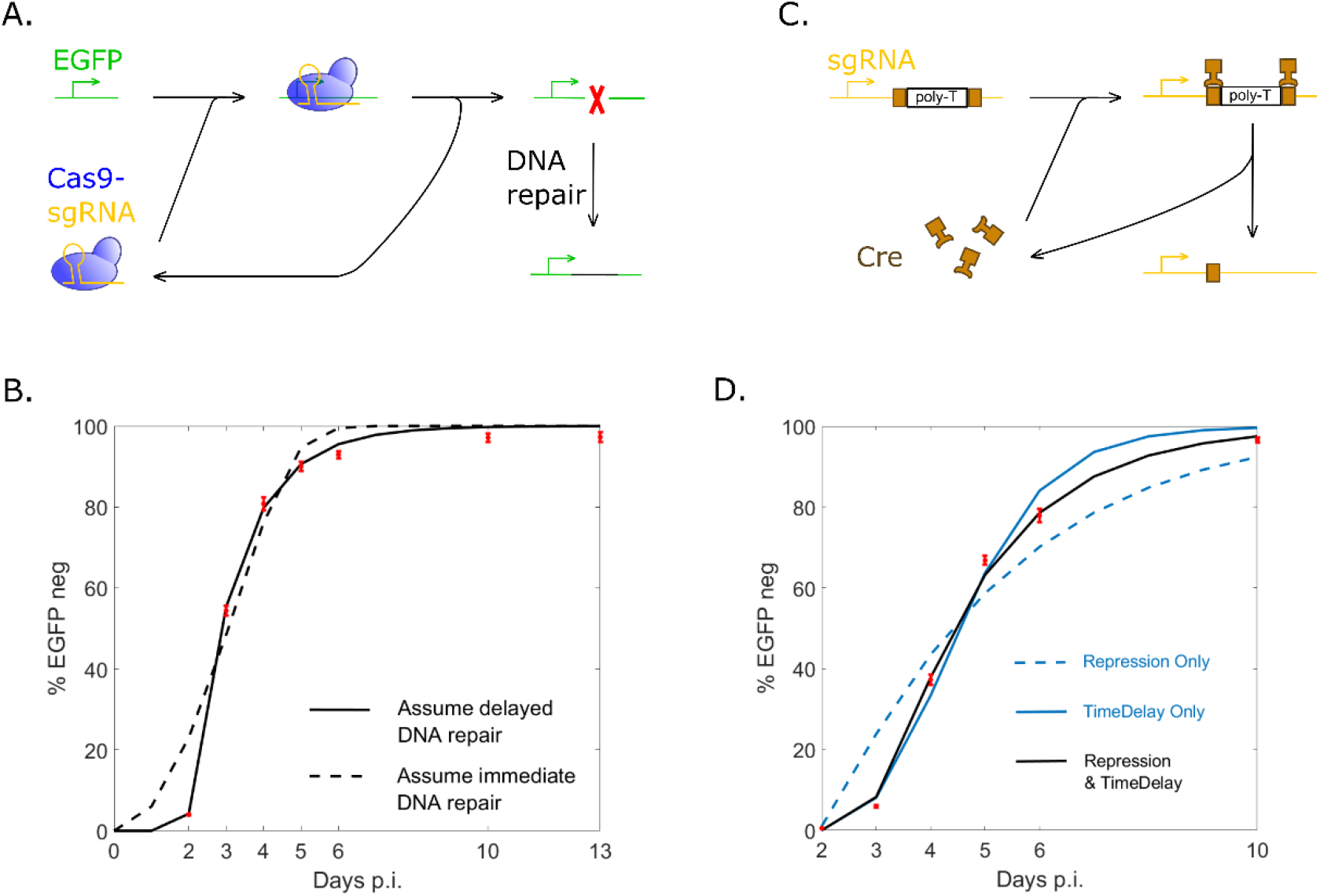
Intrinsic time delay in biomolecular activities. (A) DNA repair after CRISPR induced double strand break. (B) Data fitting for DNA repair with different assumptions. Model Assuming delayed DNA repair gives the best fit. (C) Cre recombination gives sgRNA transcription. (D) Data fitting for Cre activity with different assumptions. Both repression and time delay exist in Cre activity.

Where [EGFPm] stands for the mutated EGFP DNA, [EGFP] stands for the active EGFP DNA, lag1[EGFP] tracks EGFP concentration somewhile before. (Table 1)

The data fitting work is carried out for both models and the results are illustrated as Fig. 6B. Coefficient of determination (R^2^) is calculated to evaluate the fit. The delayed DNA repair assumption yields a R^2^ equals to 0.9981 while the immediate DNA repair assumption yields a smaller R^2^ equals to 0.9577, which supports the assumption that there’s a time delay in CRISPR induced DNA repair. The delay time for DNA repair process is estimated as 44.3 h. Fitted parameters about CRISPR activity are obtained and then used in linear safety switch (Fig. 3B, Fig. 7A), artificial time delayed safety switch (Fig. 3A) and intrinsic time delayed safety switches (Fig.7 B-D).

**FIGURE 7.**
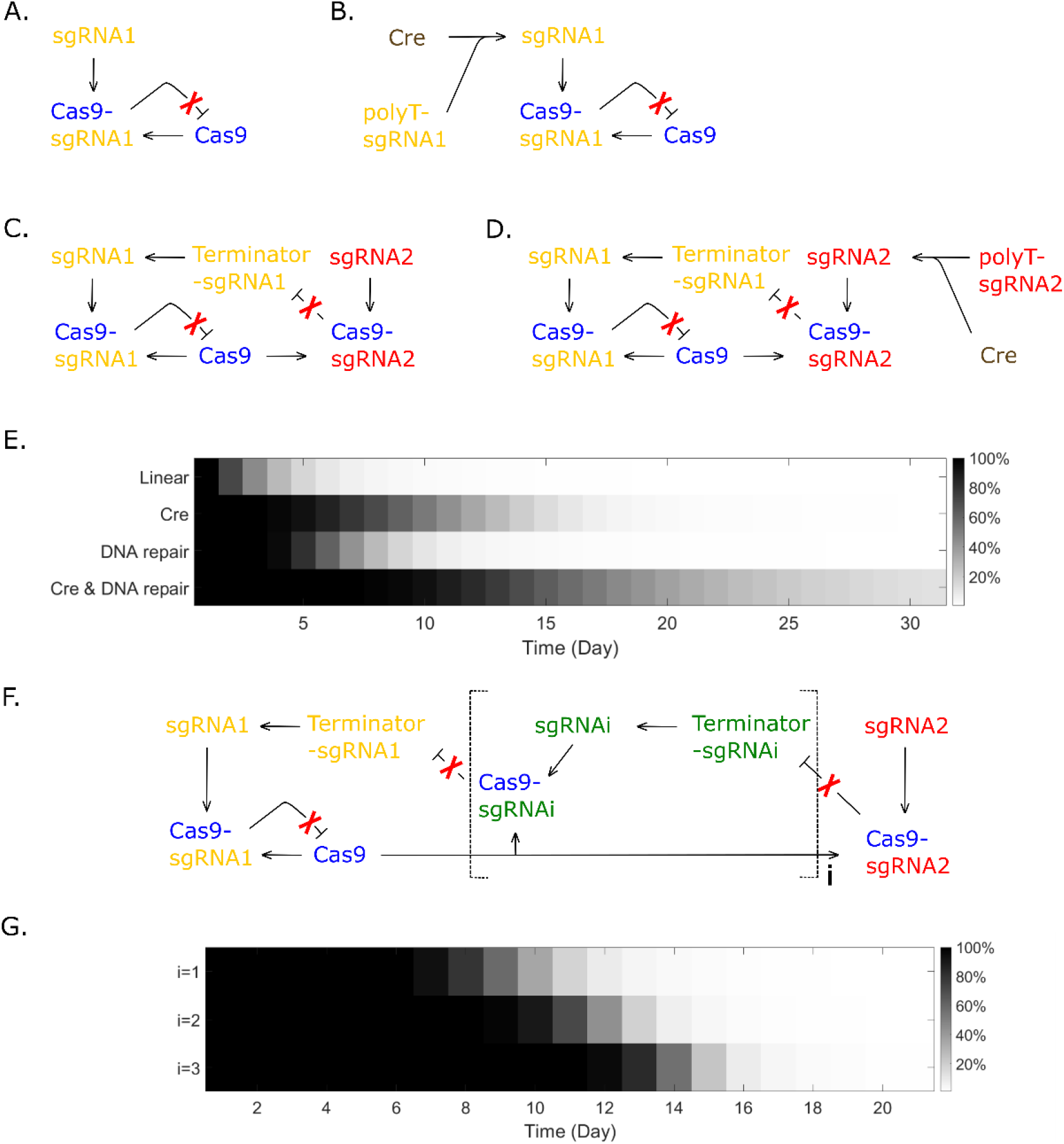
Safety switches utilizing intrinsic time delay. (A) Linear Cas9 elimination. (B) Time-delayed Cas9 elimination by Cre recombination. (C) Time-delayed Cas9 elimination by DNA repair. (D) Time-delayed Cas9 elimination by Cre recombination and DNA repair. (E) Dynamic process of Cas9 elimination for different circuits. (F) Time-delayed Cas9 elimination by cascade connection of DNA repair. (G) Dynamic process of Cas9 elimination with different number (i) of cascades.

Another circuit tested by Krzysztof et al. [30] draws our attention and it’s illustrated as Fig. 6C. In this circuit, Cre recombinase targets the two loxp sites between promoter and sgRNA coding sequence, two loxp sites are recombined into one loxp site and thereby poly-T is removed. sgRNA is transcribed after Cre recombination and induces double strand break in EGFP with Cas9. Compared to the unmodified sgRNA without loxp sites and poly-T, the sgRNA which keeps a single loxp shows a delay in EGFP mutation. It’s cited in the paper that sgRNA with a single loxp may be repressed by Cre binding activity, but the parallel pattern of delay makes us suspect that the reason could be a time delay for Cre recombination process. To evaluate different assumptions, 3 models are constructed: the first model assumes that sgRNA with loxp is repressed by Cre (equation 25), the second model assumes that there’s a time delay in Cre recombination (equation 26), and the third model assumes that both repression and time delay exists (equation 25-26).

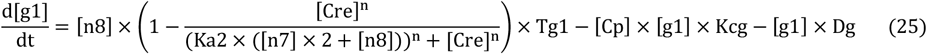

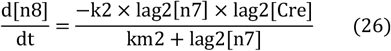

Where [Cre], [n7] and [n8] stands for Cre recombinase, DNA encodes promoter-loxp-(ploy-T)-loxp-sgRNA and DNA encodes promoter-loxp-sgRNA, respectively. Ka2 is the ratio of Cre recombinase to loxp which produces half occupation. Hill coefficient n is set to 1 because Cre is assumed to work as monomer [31]. k2 and km2 are parameters describing the activity of Cre recombination, lag2 tracks the concentration somewhile before. (Table 1)

The result of data fitting is illustrated as Fig. 6D. The repression only assumption yields a R^2^ equals to 0.9349, the time delay only assumption yields a R^2^ equals to 0.9902, while the assumption that both repression and time delay exists gives the best fit with a R^2^ equals to 0.9976. The delay time for Cre recombination process is estimated as 27.5 h. Fitted parameters about Cre activity are obtained and then used in intrinsic time delayed safety switches (Fig. 7B, Fig. 7D).

#### 3.2.2 | Safety switches utilizing intrinsic time delay

Safety switches utilizing intrinsic time delay are designed as Fig. 7B-D. To generate time delay with DNA repair, terminator flanked with sgRNA2 target site can be inserted between promoter and sgRNA1 coding sequence. Since termination efficiency is highly sensitive to the change in sequence, we expect indels generated by DNA repair will deactivate the terminator and enable sgRNA1 transcription. A self-cleaving ribozyme ahead of sgRNA1 protospacer may also be necessary in this case.

Safety switches are tested with fitted parameters and the dynamic process of Cas9 elimination is given as Fig. 7E. Safety switch with DNA repair keeps a high Cas9 plasmid concentration for 4 days and then eliminates Cas9 quickly, which shows an ideal time-delayed feature. Although the delay time estimated for Cre recombination is shorter than DNA repair, Cas9 elimination is slower in Cre recombination design since sgRNA1 transcription is repressed by Cre, the speed in self-elimination after delay time does not switch to a fast pattern. Through a combination of Cre recombination and DNA repair (Fig. 7D), Cas9 maintenance can be extended to weeks, but the pattern of Cas9 self-elimination does not have an obvious time-delayed feature. Since Safety switch using DNA repair solely has the desired time-delayed feature, we further developed this design by introducing a cascade connection (Fig. 7F) in hope that delay time can be manipulated. The result (Fig. 7G) indicates that delay time for DNA repair design is tunable by manipulating the number (i) of cascades.

## 4 | Discussion

Ultrasensitivity is a broadly discussed topic in the field of biochemistry and synthetic biology. A lot of paper about ultrasensitive cellular events [23-24], methods of generating ultrasensitivity [25-27] and circuit construction with ultrasensitivity [32-33] can be found. To the best of our knowledge, few examples of applying ultrasensitivity to therapeutic purpose are available. Here we report an ultrasensitivity transmission module which generates artificial time delay in Cas9 self-elimination. In this module, ultrasensitive relationship between Cas9-sgRNA complex (output) and Cas9 plasmid concentration (stimulus) is built. As Cas9-sgRNA complex shears Cas9 plasmid as an enzyme, an ultrasensitive relationship between enzyme and substrate naturally generates a time-delayed pattern in substrate digestion.

Our data fitting work reveals the intrinsic time delay in both DNA repair and Cre recombination but the molecular mechanism for the delay remains unknown. Compared to the tunable delay time generated by ultrasensitivity, intrinsic delay time is fixed. A cascade connection of DNA repair modules extends the delay time, but the dose of sgRNA needs to increase in an exponential pattern with the number of cascades because of the competition for limited Cas9 resource. Thus, a large number of cascade connections may not be practical in wet lab experiments. Intrinsic time delay of DNA repair and Cre recombination is reported but the existence of time delay may not be limited to these two biomolecular activities only. Discovery of time delay in other biomolecular activities may offer various delay time, less exogenous genes and other features. However, as synthetic biologists usually focus on whether their circuit works or not rather than the dynamic process, few data can be used for system biology to reveal interesting features including but not limited to time delay.

Previously reported safety switches all share the same mechanism that a sgRNA under constitutive promoter is designed to target Cas9, thereby Cas9 will be cleared. In this way, gene therapy and self-elimination occurs at the same time, and sometimes Cas9 will be cleared before gene therapy is complete. Our time-delayed safety switches separate gene therapy process and self-elimination process by introducing a time delay to self-elimination. Since the length of delay time is highly tunable, the system can balance gene therapy efficiency and side effect minimization for all kinds of clinical cases.

By reporting the time-delayed self-elimination systems which ensure both gene therapy efficiency and side effects minimization, and offering an example of reviewing data by mathematical modeling and data fitting, we hope our modeling work may provide inspiration in achieving safe and efficient gene therapy in humans.

## ACKNOWLEDGEMENTS

The author sincerely appreciates the financial support from Lixia Wang.

## CONFLICTOFINTEREST

There is no conflict of interest.

