## Supporting Information for "Design of time-delayed safety switches for CRISPR gene therapy" (from another version of this preprint: version 2, 2021-04-19)


### Dashan Sun


#### 1 | Sensitivity analysis

In our work, steady-state response curve is used to describe how the output of the system (sgRNA transcription) depends on different inputs. To quantify the degree of sensitivity, instantaneous sensitivity [1] is defined as the equation below:

$$s(I) = \frac{\mathrm{dO/O}}{\mathrm{dI/I}} = \frac{\mathrm{dlog(O)}}{\mathrm{dlog(I)}}$$

Where O stands for output and I stands for input. Response is ultrasensitive if |s| > 1, linear if |s| = 1, subsensitive if 0 < |s| < 1 and desensitive if s = 0.

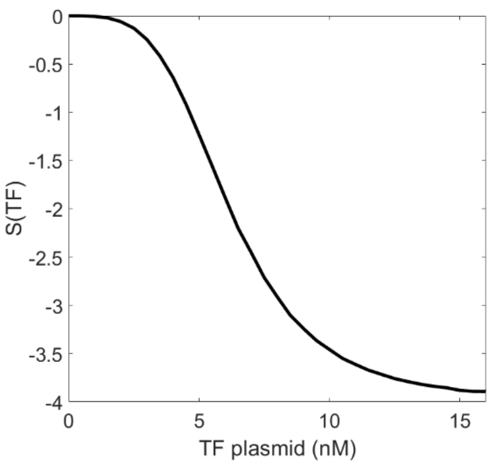


**F I G U R E S1** Sensitivity analysis for sgRNA (output) and transcription factor plasmid (input). TF stands for transcription factor.

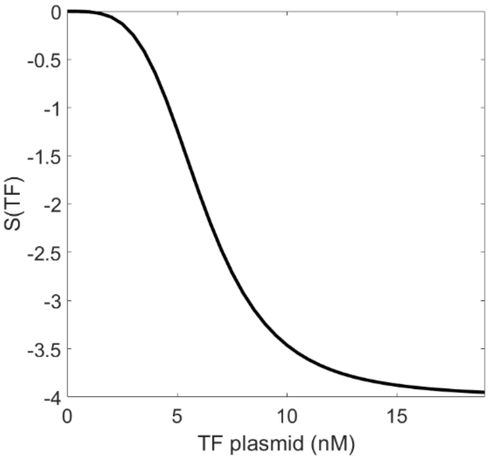


**F I G U R E S2** Sensitivity analysis for Cas9-sgRNA (output) and transcription factor plasmid (input).

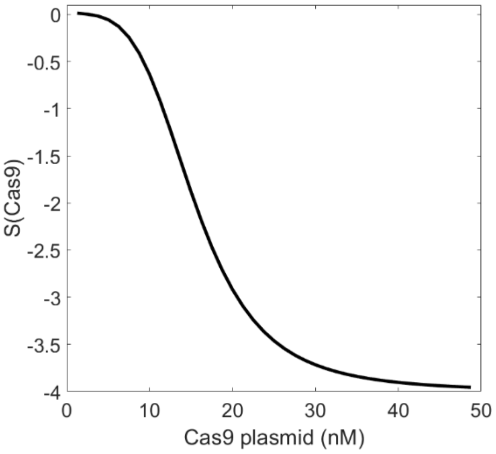


**F I G U R E S3** Sensitivity analysis for Cas9-sgRNA (output) and Cas9 plasmid (input).

### 2 | Cas9 activity in multiple target design

The process of Cas9-sgRNA shearing plasmids is modeled as enzyme catalyzing substrates. Specifically, Cas9-sgRNA complex is enzyme (E), promoters inserted with gRNA targeting site are substrates (S), sheared promoters are products (P) of reactions. In our work, the same Cas9-sgRNA targets different promoters and reactions are described as the equation below.

$$Sn + E \underset{k-1}{\overset{k1}{\rightleftharpoons}} E{:}Sn \xrightarrow{k2} E + Pn$$

According to Quasi-steady-state approximation,

$$\frac{d(E:Sn)}{dt} = 0 \quad n = 1, 2, \dots, m \quad (1)$$

Thus,

$$E:Sn = \frac{k1 \times Sn \times E}{(k-1) + k2} \quad n = 1, 2, \dots, m \quad (2)$$

According to enzyme conservation,

$$Et = E + E:S1 + E:S2 + \cdots + E:Sm \quad (3)$$

Speed of Sn elimination is described as

$$\frac{dPn}{dt} = k2 \times E:Sn \quad n = 1,2, \dots, m \quad (4)$$

Combine equations 2, 3, 4, we get:

$$\frac{dPn}{dt} = \frac{k2 \times Sn \times Et}{km + S1 + S2 + \cdots + Sm} \quad (5)$$

Where km=(k-1+k2)/k1.

### 3 | Data fitting

We have the parameters fitted in a range and frequency by running the lsqcurvefit function with random initial values for 40 times. Parameters for Cas9 activity and Cre recombination are fitted as Fig. S4 and Fig. S5, respectively.

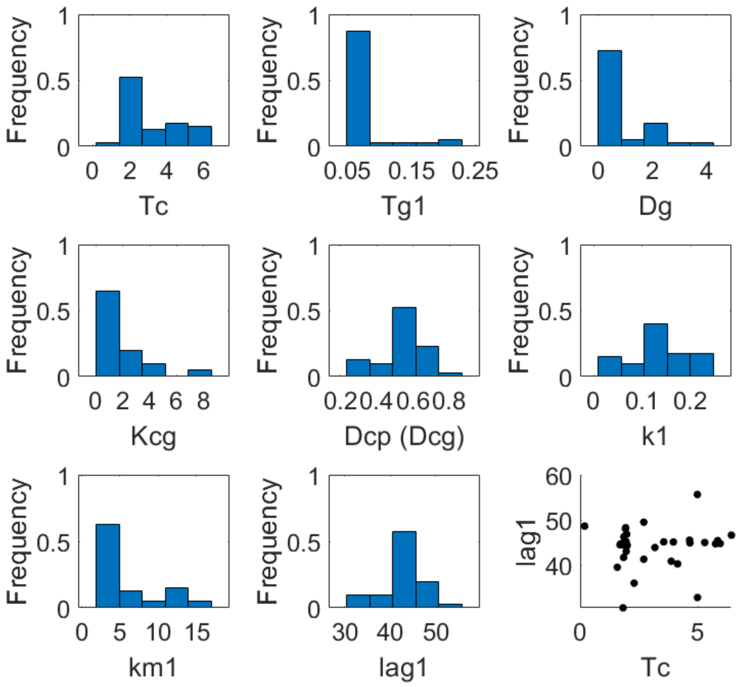


**F I G U R E S4** Parameters for Cas9 activity.

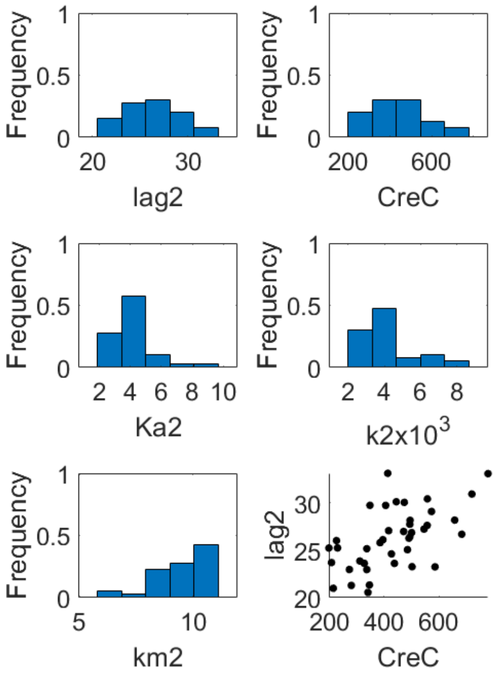


**F I G U R E S5** Parameters for Cre recombination.

### 4 | Parameters

**TABLE S1** Parameters

| Parameters | Descriptions | Value | Reference |
|---|---|---|---|
| [n1] – [n5] | Plasmid concentration | 5nM, 46nM, 50nM | Estimated |
| [n6] – [n8] | Concentration of ORF in genome | 10nM | Estimated |
| T1 | Transcription factor transcription rate | 0.6 (/h) | [2-3] |
| T2 | Transcription factor translation rate | 1.4 (/h) | [4] |
| Tg | sgRNA transcription rate | 0.06807 (/h) | Data fitting |
| Tcr | Cas9 transcription rate | 4 (/h) | Estimated |
| Tcp | Cas9 translation rate | 0.6 (/h) | [4] |
| Tc | Cas9 protein generation rate | 2.6514 (/h) | Data fitting |
| Dtfr | Transcription factor mRNA degradation rate | 0.06 (/h) | [6] |
| Dtfp | Transcription factor protein degradation rate (with degradation tag) | 0.6 (/h) | Estimated |
| Dg | sgRNA degradation rate | 1.15138 (/h) | Data fitting |
| Dcr | Cas9 mRNA degradation rate | 0.06 (/h) | [6] |
| Dcp, Dcg | Cas9 protein degradation rate, Cas9-sgRNA complex degradation rate | 0.2557 (/h) | Data fitting |
| Ka1 | Transcription factor concentration producing half occupation to 1 nM target | 3 (nM) | Estimated |
| Ka2 | Cre recombinase producing half occupation to 1 nM loxp site | 4.2228 (nM) | Data fitting |
| Kcg | Cas9 and sgRNA binding rate | 0.3119 (/(nM*h)) | Data fitting |
| k1 | Parameter about Cas9-sgRNA shearing target DNA | 0.1308 (/h) | Data fitting |
| km1 | Parameter about Cas9-sgRNA shearing target DNA | 11.7129 (nM) | Data fitting |
| k2 | Parameter about Cre recombination | 0.004 (/h) | Data fitting |
| km2 | Parameter about Cre recombination | 12.7710 (nM) | Data fitting |
| d | Dilution rate | 0-0.1 (/h) | [5] |
| n | Hill coefficient | 1-6 | Estimated |
| lag1 | Delay time for DNA repair | 44.2974 (h) | Data fitting |
| lag2 | Delay time for Cre recombination | 27.4716 (h) | Data fitting |
| CreC | Cre concentration in steady state | 489.1786 (nM) | Data fitting |
